# Examining Batch Effect in Histopathology as a Distributionally Robust Optimization Problem

**DOI:** 10.1101/2021.09.14.460365

**Authors:** Surya Narayanan Hari, Jackson Nyman, Nicita Mehta, Haitham Elmarakeby, Bowen Jiang, Felix Dietlein, Jacob Rosenthal, Eshna Sengupta, Alexander Chowdhury, Renato Umeton, Eliezer M. Van Allen

## Abstract

Computer vision (CV) approaches applied to digital pathology have informed biological discovery and development of tools to help inform clinical decision-making. However, batch effects in the images have the potential to introduce spurious confounders and represent a major challenge to effective analysis and interpretation of these data. Standard methods to circumvent learning such confounders include (i) application of image augmentation techniques and (ii) examination of the learning process by evaluating through external validation (e.g., unseen data coming from a comparable dataset collected at another hospital). Here, we show that the source site of a histopathology slide can be learned from the image using CV algorithms in spite of image augmentation, and we explore these source site predictions using interpretability tools. A CV model trained using Empirical Risk Minimization (ERM) risks learning this source-site signal as a spurious correlate in the weak-label regime, which we abate by using a training method with abstention. We find that a patch based classifier trained using abstention outperformed a model trained using ERM by 9.9, 10 and 19.4% F1 in the binary classification tasks of identifying tumor versus normal tissue in lung adenocarcinoma, Gleason score in prostate adenocarcinoma, and tumor tissue grade in clear cell renal cell carcinoma, respectively, at the expense of up to 80% coverage (defined as the percent of tiles not abstained on by the model). Further, by examining the areas abstained by the model, we find that the model trained using abstention is more robust to heterogeneity, artifacts and spurious correlates in the tissue. Thus, a method trained with abstention may offer novel insights into relevant areas of the tissue contributing to a particular phenotype. Together, we suggest using data augmentation methods that help mitigate a digital pathology model’s reliance on potentially spurious visual features, as well as selecting models that can identify features truly relevant for translational discovery and clinical decision support.

## 1. Introduction

Computer vision (CV) approaches applied to cancer histopathology image data have demonstrated emerging potential for biological discovery, precision diagnostics, and as predictive biomarkers [1, 2, 3, 4, 5]. However, challenges persist regarding the computational, interpretability and generalization realms stemming from the giga-pixel nature of the Whole Slide-Images (WSIs) and the absence of patch-level labels. Previous efforts [6, 7] attempt to address the computational and interpretability challenges. However, generalizability is still an unsolved challenge that could result in variable performance among underrepresented subpopulations of patients in each hospital [8, 9, 10, 11]. These generalizability issues have resulted in considerable decision-making complexity when implementing solutions using deep learning in digital pathology.

### 1.1. Spurious confounders in digital pathology

For CV applications, lack of model generalizability is often a result of the effect of spurious correlates introduced as a result of the WSI preparation process, also known as batch effects [12, 13, 14]. Mitigating all forms of batch effects parametrically incurs challenges since batch effects may arise from different parts of the tissue preprocessing pipeline such as the scanner acquisition protocol, slide preparation date and thickness of tissue sections [15, 16, 17, 18]. These batch effects remain detectable by machine learning algorithms and can induce spurious correlates. Methods have been proposed to solve visible batch effects [19, 20]. However, such methods cannot fully account for subtle batch effects that might persist, such as distinct patient demographic profiles in different hospitals that result in different biological and clinical baseline features specific to data derived from each hospital. Indeed, multiple studies have demonstrated that a trained model can learn the race and age of a patient [15, 17], and these features might also serve as confounders to the model.

### 1.2. Distributionally Robust Optimization (DRO) as a solution to prevent learning spurious confounders

When patch-level models that are trained using slide-level labels exhibit a low training error, they might have done so by learning spurious correlates from the patches that do not exhibit features of the slide-level label. This potential overfitting to spurious correlates could exacerbate disparities that exist due to underlying differences in patient populations served at different hospitals, among other factors [21, 22]. For example, if all tumors of Lung adenocarcinoma (a subtype of lung carcinoma, the other major subtype being squamous cell carcinoma) are all higher grade in the training set, but lower grade in the validation set, and vice versa for squamous cell carcinoma, we want a model to be robust to the distributional shift in the grade between the training and validation sets, while performing the subtyping task. A subfield of DRO, Group-DRO [21], aims to increase robustness to shifts in the groups between training and validation sets. However, this approach requires expert annotation to explicitly characterize and enumerate the groups of the cancer tissues.

In addition, when a digital pathology model is trained on data from one source hospital and tested on data from the same hospital, it could over-fit to batch effects instead of fitting to an outcome-wide distribution that generalizes to other source sites. This problem is often abated by having an external test set [10, 23, 24, 25, 26, 27], since using independent methods of data-collection helps validate generalization. The task then generalizes to features that can be observed in a variety of settings with different preprocessing methods. However, testing on a diverse held-out set requires holding out data from the training process and deprives the training process of this diversity.

### 1.3. Evaluating proposed solution on tasks with clinical relevance

Existing solutions to circumvent the potential confounding introduced by the spurious correlates include methods to resolve intra-WSI heterogeneity [28]. However, these methods include computational overheads and more hyperparameters. To circumvent this, we propose training using an abstention method. Here, we evaluate using a groupDRO method and a model trained with abstention relative to established approaches across three CV histopathology tasks with clinical relevance:

#### 1.3.1 Lung Carcinoma

Lung adenocarcinoma (LUAD) is one of the two major histologic subtypes of Non-Small Cell Lung Cancers (NSCLC), the other being Lung Squamous Cell Carcinoma (LUSC). LUAD and LUSC affect nearly 40% and 20% of all Lung cancer patients in the United States [29]. Identification and subtyping of the tumor in a WSI can help guide pathologic assessment, as well as potentially determine the efficacy of therapy [2, 30, 31]. However, identification of tumor may be confounded by scarring tissue from the effects of smoking on lung tissue, amongst other features.

#### 1.3.2 Predicting grade in clear cell Renal Cell Carcinoma

Clear cell Renal Cell Carcinoma (ccRCC) makes up 80% of the incidence of all Kidney Cancer cases, which will affect an estimated 76,000 people in the United States during 2021 [32, 33]. In patients with ccRCC, amongst pathological features classified based on cell shape and arrangement, nuclear size, nuclear irregularity and nucleolar prominence showed highest effectiveness in predicting distant metastasis, even more so than tumor size [34]. These features are used to grade the tumor, with a higher grade implying worse prognosis. These morphological features can be distinguished visually and offer potential for the application of CV algorithms. However, due to inter-observer variability and intra-tumoral heterogeneity, CV algorithms are susceptible to batch effects and confounding by spurious correlates.

#### 1.3.3 Predicting Gleason score in prostate adenocarcinoma

Prostate adenocarcinoma (PRAD) forms the large majority of prostate cancers, which will affect just shy of a quarter of a million people in the United States during 2021 [35, 36]. The Gleason grading system tailored to specific properties of this histology is used to describe the patterns observed in tumor tissue in prostate adenocarcinoma (PRAD), with grades ranging from 1 (least advanced) to 5 (most advanced). A Gleason score for the sample biopsy is then calculated by adding the two most prominent grades visible in the tissue. In practice, the lowest Gleason score awarded that qualifies as cancer is a 6 (3+3). Recent works have shown the use of CV to predict the Gleason score of a scan of biopsy tissue [37, 38]. However, whether or not Gleason scoring models are learning spurious correlates of the Gleason grade is incompletely characterized but critical for clinical use.

## 2. Methods

Here, we propose a new training method (henceforth referred to as training with *abstention*) that we compare with conventional backpropagation using Empirical Risk Minimization (ERM). A full overview of our pre-processing pipeline is elaborated in figure 1a. Gigapixel whole slide images (WSIs) are first passed through a quality control (QC) process using HistoQC [39] and subsequently divided into a number of patches (order of 10^3^) in a process called *tiling*. Subsequently, they are first augmented through a color jitter or stain normalization process, described below, after which, they are passed through machine learning models described in section 2.1 onward.

**Figure 1:**
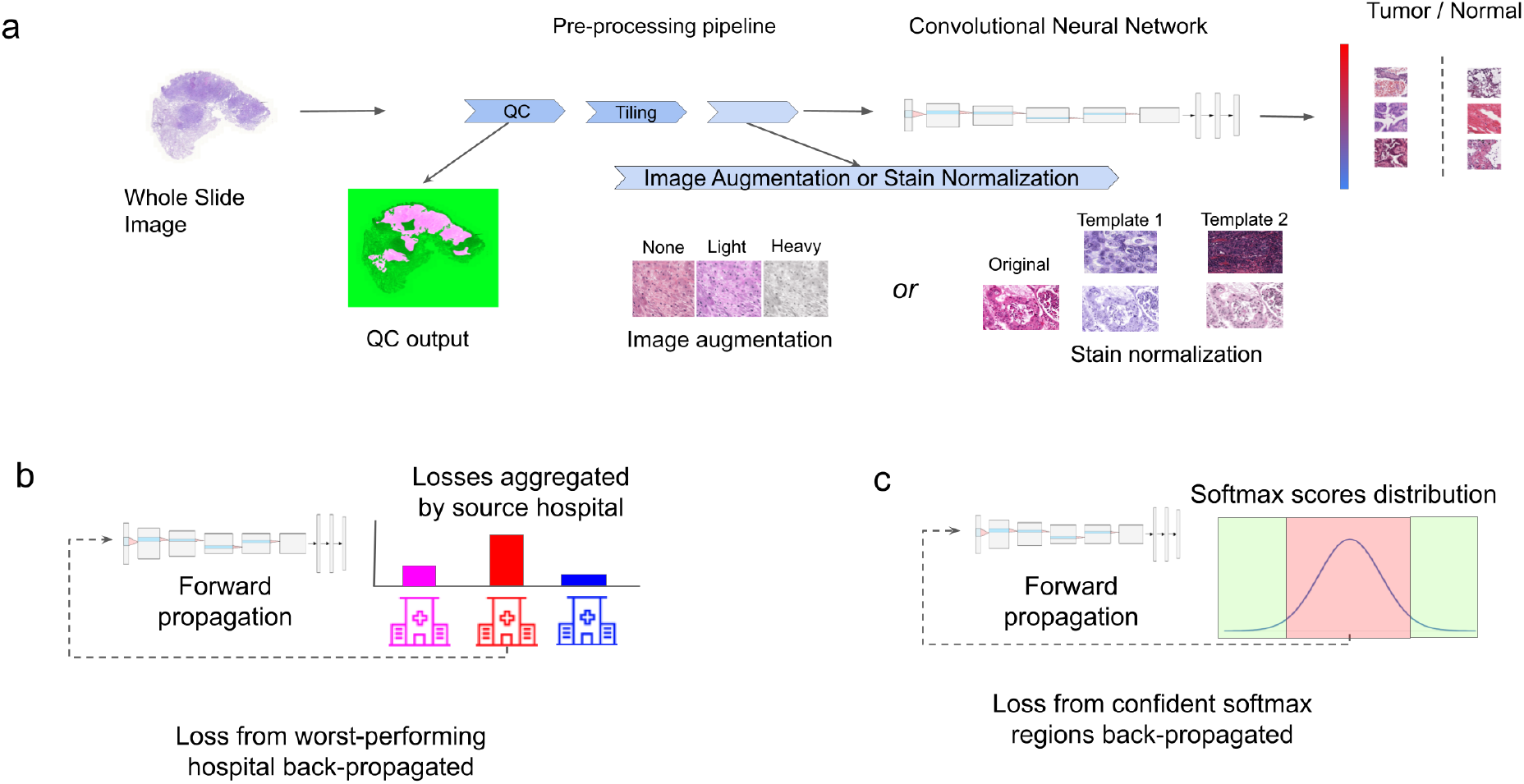
Models used in our experiments, a) showing a standard ERM model used in our pipeline to predict whether a patch comes from tumor tissue or surrounding healthy tissue b) a group-DRO algorithm that defines groups based on the source hospital of the patch c) An algorithm that updates its weights based on loss accumulated on examples that it is confident on

### Color Jitter

We used image augmentation via jittering the RGB pixel values in the RGB space to prevent overfitting to the color distribution by inducing random changes in the brightness, saturation, and other properties of an image, also known as color jitter [40]. To discretize the color jitter, we defined a light version of the color jitter that allowed the brightness factor to be chosen uniformly at random between [0.875, 1.125], the contrast factor to be chosen uniformly at random between [0.5, 1.5], the saturation factor to be chosen uniformly at random between [0.5, 1.5] and the hue factor to be chosen between -0.1 and 0.1. We similarly defined a heavy version of the color jitter to be four times proportionally higher (unless limited by the maximum allowed limits for each factor). That is, we allowed the brightness factor to be chosen uniformly at random between [0.5, 1.5], the contrast factor to be chosen uniformly at random between [0, 3], the saturation factor to be chosen uniformly at random between [0, 3] and the hue factor to be chosen between [-0.4, 0.4]. The limit on the color jitter we could introduce was placed by the hue factor, which was forced to be between [-0.5, 0.5].

### Stain Normalization

We also used image augmentation via jittering the RGB pixel values in the RGB space to prevent overfitting to the color distribution. In addition to using color augmentation, we also used stain normalization using Staintools [41]. We performed stain normalization in two ways: 1) Where the images in the validation set were normalized to the same template as the images in the training set and 2) Where the images in the validation set were normalized to a different template compared to the images in the training set. The first method was used to prevent the stain template of the image from creating a spurious correlate. The second method was used to test the model’s reliance on morphological features that are still observable despite a distributional shift in the color profile. However, we did not use stain normalization in our tasks with clinical relevance owing to the performance bottleneck imposed by the stain normalization process.

### 2.1. ERM model

In order to establish a baseline to compare our models trained with group distributionally robust optimization (group-DRO) and trained with abstention, we use a pretrained ResNet-50 convolutional neural network (CNN) [42]. The model was pretrained on the ImageNet dataset [43]. We replaced the final layer with a layer having a number of heads pertaining to the number of classes in our task whose weights are initialized uniformly at random [44]. We used a cross-entropy loss function where the loss is computed and aggregated over the entire dataset. This model is henceforth referred to as the ERM model.

### 2.2. group-DRO

In our implementation of a group-DRO method, we defined the groups as hospitals from which the WSIs were taken. We trained an algorithm by backpropagating the loss over the tiles from the worst performing hospital, measured by average loss per tile in a batch. However, the reported statistics, such as F1, are reported over the whole validation / testing dataset, and not the worst performing hospital. A pictorial representation of this algorithm is shown in Figure 1b.

#### Algorithm 1

Forward Propagation of Loss in Abstention architecture

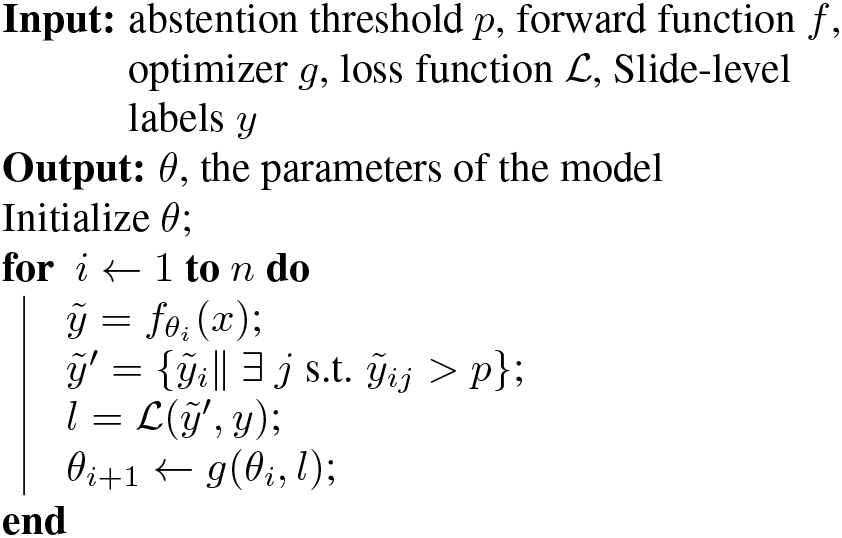

### 2.3. DRO with abstention

Models were trained using an abstention algorithm (Algorithm 1, Figure 1c) whereby we only accumulated and backpropagated the losses from images for which the model predicted a maximum softmax logit score greater than a predefined threshold, *p*. We interpret this threshold as a confidence and only report losses on images for which the confidence value is greater than *p*. We used this abstention method while training so that the weights learned by the model are on data that the model is confident about. To rescale the outputs of the softmax function into a probability distribution for thresholding by *p*, we used temperature scaling [45].

### 2.4. Training details

#### Cross Validation

We performed 5-fold cross validation. We allowed folds to overlap with one-another at the slide or patient level depending on the task. In the set of experiments where we trained on one hospital and evaluated on another, we only performed three cross validation trials.

#### Early Stopping

We train our models to minimize error and stop training if the error does not improve on the validation set over five consecutive measurements [46]. The validation performance was measured four times per epoch. Thus, a lack of performance improvement for five consecutive measurements implies that the model’s validation performance did not increase over one epoch. We found the patience of five to be suitable through cross validation performance.

#### Reporting F1

We reported the best validation F1 achieved by the model, unless stated otherwise. We continued to track the loss metric to evaluate further improvement by the model; however, an improvement in loss does not necessarily improve F1. Thus, we report the F1 at the training instant where the F1 is highest even if the model achieves a lower loss at a different time point.

## 3. Experiments

### 3.1. Predicting the source site of a histopathology tissue

We used an ERM model to predict a scan’s source hospital for LUAD patches. We trained the model with a one-hot encoded label of the source site of the image as the label.

#### Data Imbalance

There was an uneven distribution of tiles across hospitals donating to TCGA. Balancing the number of WSIs and the number of QC-checked tiles from each hospital proved challenging as some hospitals contributed only a single WSI. Thus, we limited our study to the ten most populous hospitals, as measured by the number of WSIs from the site.

#### Data Splitting

The data were split into training and validation sets using data from held out patients.

#### Interpretability

We leveraged Grad-CAM [47] as an initial step in interpretability. Grad-CAM produces arrays with the same shape as the input image, which can be overlaid over the image to produce heatmaps.

### 3.2. Comparing ERM vs. DRO

We compare our method of training with abstention against ERM methods in three classification tasks with clinical relevance. We provide more relevant details on the tasks below.

#### 3.2.1 Lung Carcinoma

In one set of experiments, We evaluated our method on the task of detecting tumor tissue in Lung adenocarcinoma (LUAD), using LUAD WSIs from the Cancer Genome Atlas (TCGA) (*n* = 522). We trained a binary patch-level classifier using slide-level labels to classify tissue patches into tumor or normal tissue.

In one set of experiments done on TCGA-LUAD, we trained the model on data taken from one hospital and validated it on data taken from another without fine-tuning, to mimic a real-world setting where data is private and cannot be shared between institutions in a resource scarce setting. In order to study the effect of the preprocessing steps employed by a singular hospital, we were limited in our analysis to data from hospitals that have both tumor samples and surrounding normal tissue.

In another set of experiments, we used a private dataset to distinguish between the two major subtypes of lung cancer cases, LUAD and Lung Squamous Cell Carcinoma (LUSC). Similar to the case of detecting LUAD, we trained a binary classifier at the patch level.

#### 3.2.2 Predicting Grade of tissue in TCGA-ccRCC

We classified patches of tumor tissues taken from TCGA-ccRCC (*n* = 504) into Grade II or Grade IV cancer using slide-level labels. In order to prevent introducing confounders to the model, we first trained a model to extract tumor tissue from the WSI. This model was trained on a task of distinguishing tumor tissue from normal tissue using pixel-level labels from an in-house dataset. We proceeded with subsequent analysis of determining the grade on patches of the WSI that showed higher likelihood of being tumor tissue than healthy tissue, as predicted by this model. We also repeated the experiments on the whole dataset without removing non-tumor tiles for the sake of completeness, with data split into training and validation using data from held-out hospitals without bleeding data from the same slide or hospital from training into validation.

#### 3.2.3 Prostate adenocarcinoma (PRAD)

We predicted the aggregate Gleason score at the patch-level of a WSI taken from TCGA-PRAD (*n* = 371) using a binary classifier of low (score of =6) or high (≥ 8). We first eliminated tiles that had a less than random chance of being tumor using predictions made on patch-wise labels and data from Schömig-Markiefka et al. [18]. We also repeated the experiments on the whole dataset without removing nontumor tiles for the sake of completeness, with data split into training and validation without bleeding data from the same slide or hospital from training into validation, unless mentioned otherwise.

#### 3.2.4 Data splitting and Training details

In the tasks on PRAD, we used a crop size of 512 and a batch size of 32. In the tasks on ccRCC and LUAD, we use a crop size of 224 and a batch size of 128. This was decided based on cross-validation experiments.

In the tasks on LUAD and PRAD, we combined the data from all hospitals, and compared models trained using DRO against models trained using ERM, and ablated the number of hospitals held out during testing, measuring the robustness of the model when more hospitals are held out. In LUAD and PRAD, when *j* hospitals held out, we took *j* held out hospitals from each class. In ccRCC, however, owing to data availability constraints, we did not ablate the number of hospitals. We instead created a validation set with roughly 30% of the unique hospitals from the entire dataset, and validated a model on data from those held out hospitals. The reported statistics (performance, coverage and errors of the same) are reported on this held out set.

For the method trained with abstention, we report test statistics, such as F1 and loss, on images for which the model reports softmax logits with confidence values greater than *p*. We compute a macro-F1, aggregating the F1 scores of the individual classes without weighting them by the number of samples.

## 4. Results

### 4.1. Heterogeneity in predicting tumor vs. normal tissue

First, we evaluated models trained on a single source site and validated on either the same or different single source site on a task of LUAD identification. Overall, we found significant heterogeneity in model performance based on the hospital whose data were used to train and validate the model (Figure 2b). For example, a model trained on data from the University of Pittsburgh achieved a validation F1 of 0.97 when validated on data from a held-out set of patients from the University of Pittsburgh, but, at best, only achieved a validation F1 of 0.71, when evaluated on data from Prince Charles Hospital.

**Figure 2:**
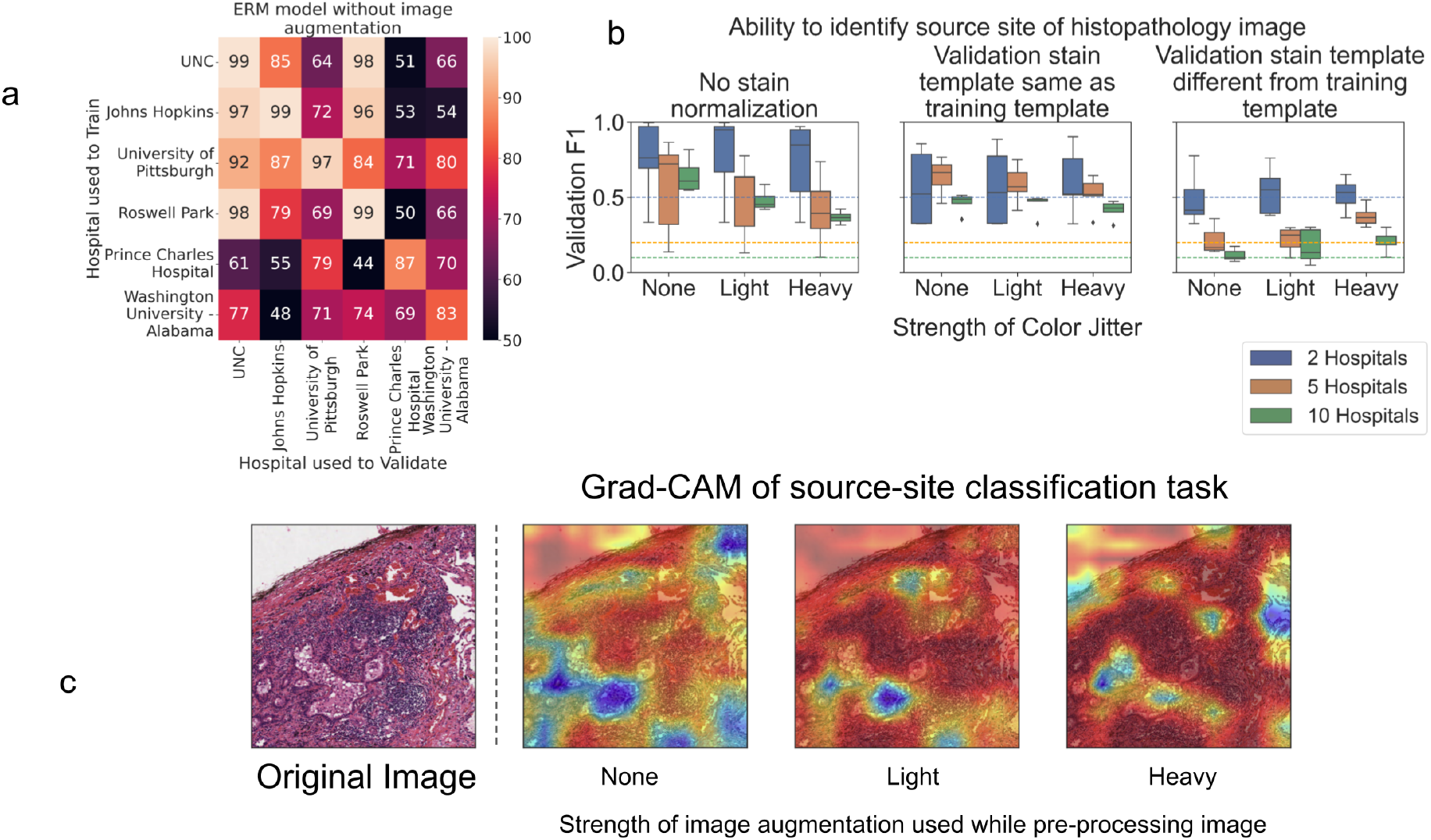
a) Heterogeneity in model performance in task to identify whether a patch contains tumor in TCGA-LUAD b) an ERM model’s performance in identifying the source site of a patch from a TCGA-LUAD WSI with image augmentation techniques applied to mask out the effect of source site c) pixel importance of source-site prediction task using Grad-CAM

We then consolidated the data by aggregating across hospitals whose data were used to train and validate, again observing inter-hospital validation heterogeneity. We also found that hospitals whose data on which models achieve a higher validation F1, did not achieve comparable performance when models trained on that same site’s data were validated on other hospitals, and vice versa. For example, a model trained on data from the University of Pittsburgh, achieved a median validation F1 of 0.86 when validated on other hospitals. However, models trained on data from other hospital sites and validated on the University of Pittsburgh cohort achieved a median F1 of 0.72. Further, for data from the hospitals at the University of North Carolina and Roswell Park, models achieved higher performance when used for validation (0.95 and 0.91 median F1, respectively) rather than for training (0.76 and 0.74 median F1, respectively).

### 4.2. Impact of image augmentation on identifying the source site of an image

Given the heterogeneity in model performance, we next evaluated a possible source of this heterogeneity that arises from the data preparation and pre-processing steps. Consistent with prior reports [48], we found that a model could recognize the source site of a histopathology scan through visual features in the absence of stain normalization (Figure 2a, left). However, we found that color jitter was able to mitigate the ability of the model to discern the source site of an image by up to 10%, 33% and 24% when distinguishing between 2, 5 and 10 hospitals respectively. To attempt to mask out the stain profile, we normalized the stain across the images. However, in spite of stain normalization, we were still able to distinguish the source hospital of an image with 67% and 49% F1 for 5 and 10 source hospitals respectively, when no color jitter was used (Figure 2b, middle). When heavy color jitter was applied, this performance decreased by 15% and 6% F1 for 5 and 10 hospitals respectively. With stain normalization such that the validation template was different from the stain template (Figure 2b, right), we found that the model performed randomly when there was no color jitter introduced, and as the color jitter strength increased to heavy, the model performance increased by 20 and 14% F1 for 5 and 10 hospitals respectively. Thus, source hospital information is, at least, in part encoded in the stain profile of the scan, which can only be partially occluded by image augmentation techniques, such as color jitter and stain normalization.

#### 4.2.1 Using Grad-CAM to identify features contributing to source-site prediction

In order to understand the features contributing to source-site signal, we used Grad-CAM [47]. We applied GradCAM to our models at various resolutions to examine the highlighted features. At both the 5x and the 20x resolutions, we found that the image augmentations did not drastically alter the regions of the image highlighted by Grad-CAM (example patch at 5x shown Figure 2c). Further, we found that Grad-CAM segmentations did not agree with any discernible boundaries of objects in the image, both at the 5x and 20x resolutions, making the masks hard to interpret. Thus, we could not identify interpretable features that contributed to source site prediction.

### 4.3. Lung Carcinoma

Given the multiple challenges presented by batch effects, we trained a model with group-DRO to evaluate whether this approach was robust to spurious confounders. When trained on data from multiple hospitals on the task to detect LUAD, we found that a model trained using groupDRO performed competitively to an ERM model, while a model with abstention outperformed a model trained using ERM under all numbers of hospitals held out, with up to 9.9% gain in F1 at the expense of 45.2% coverage (Table 1). Thus, application of DRO and group-DRO methods for the task of identifying tumor tissue in LUAD showed promise for broader applicability.

**Table 1:** Comparing proposed models against an ERM model in the case of identifying tumor tissue vs. surrounding benign tissue in LUAD

| # of Hosp. held out | ERM | Best DRO Model |  |  | group-DRO |
| --- | --- | --- | --- | --- | --- |
|  | F1 | F1 | Threshold | Coverage | F1 |
| 1 | $91.1 \pm 4.05$ | $98.4 \pm 7.89$ | 0.8 | $86.6 \pm 5.80$ | $92.2 \pm 8.08$ |
| 2 | $88.3 \pm 2.29$ | $95.2 \pm 4.39$ | 0.9 | $54.3 \pm 15.0$ | $88.6 \pm 3.99$ |
| 3 | $78.6 \pm 8.78$ | $82.0 \pm 9.26$ | 0.9 | $65.2 \pm 20.8$ | $83.6 \pm 9.84$ |
| 4 | $81.1 \pm 8.86$ | $91.0 \pm 7.70$ | 0.9 | $54.8 \pm 21.8$ | $88.8 \pm 5.85$ |
| 5 | $72.2 \pm 3.84$ | $79.7 \pm 7.37$ | 0.9 | $59.6 \pm 23.8$ | $77.4 \pm 5.09$ |

In the set of experiments where we trained a model on data from one hospital and measured its performance on data from another, we also found that heavy color jitter produced only up to 0.15 improvement in F1 and using our abstention model produced up to 0.24 improvement in F1 when used in conjunction with heavy color jitter (Figure 3a). To this effect, we propose using the DRO model to be more robust to the heterogeneity in training data and OOD validation data.

**Figure 3:**
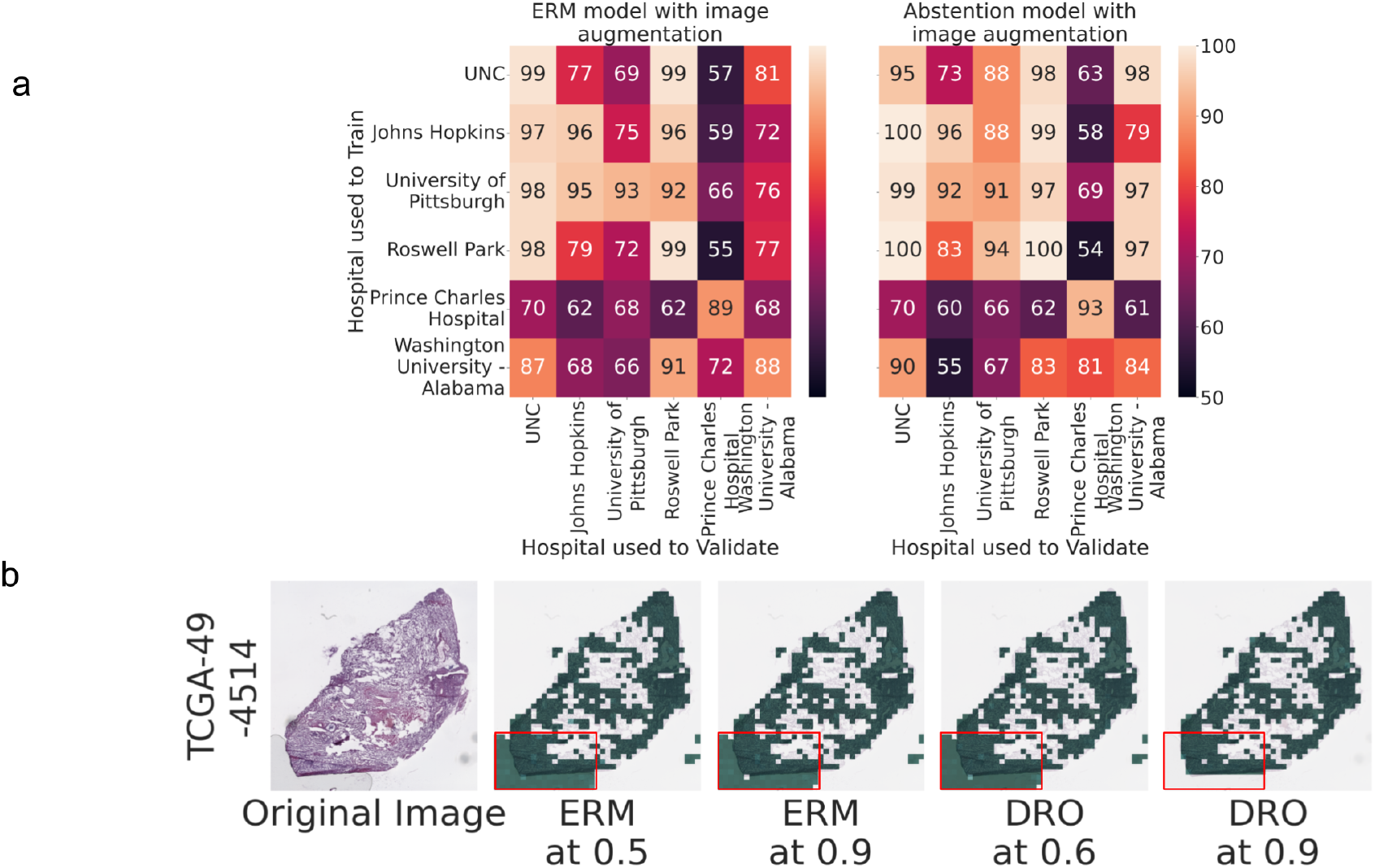
a) Image augmentation improves heterogeneity in performance in task of identifying tumor tissue in TCGA-LUAD (left) and training by abstention further improves heterogeneity (right). b) Qualitative examination shows model trained with abstention at high thresholds abstains from making histological predictions on spurious correlates of air bubbles

Upon investigation, we noted that these methods abstained from making predictions on regions of the WSI covered by slide-preparation artifacts, such as air bubbles (Figure 3b), making it less likely to learn spurious correlates.

Further, we found that models trained with abstention also abstained from more biological spurious correlates (Figure 4). In an example taken from a brain biopsy of a metastatic lung cancer, we observed i) an ERM model placed importance on surrounding brain tissue which was confirmed by a pathologist to not bear any tumor, and thus had learned spurious signal; and ii) models trained with stringent abstention in contrast completely disregarded the brain tissue, while placing modest confidence in the verified lung tumor tissue. This example was at full coverage, where all tiles of Whole Slide Image (WSI) are shown. However, owing to the different training processes, the models learned different features.

**Figure 4:**
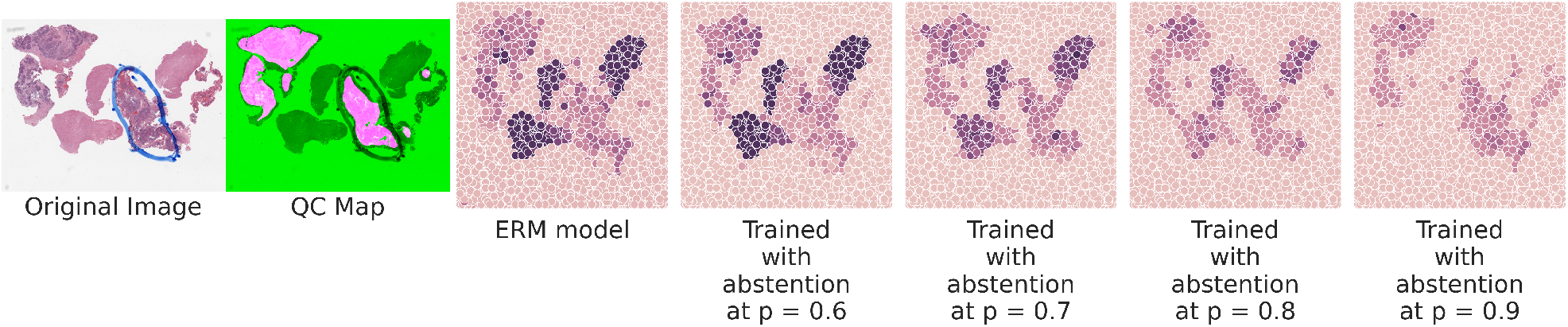
Qualitative examination shows model trained with abstention at high thresholds abstains from making histological predictions on spurious correlates of surrounding brain tissue in the case of subtyping metastatic lung carcinomas

To further demonstrate the differences in features learned in each model type, we used each model to separately produce “pruned” datasets at varying degrees of confidence, and then used these datasets to train a further set of ERM models to distinguish lung subtypes. At higher confidence levels (0.8 and 0.9), models trained on DRO-pruned data offered better performance than those trained on ERM-pruned data (0.61 ± 0.12 vs. 0.42 ±0.11 F1 [*p* = 0.10] at threshold 0.8; 0.81 ± 0.12 vs 0.53 ± 0.12 F1 [*p* = 0.047] at threshold 0.9).

### 4.4. Using group-DRO to improve generalization in grade prediction in TCGA cc-RCC

Regarding cc-RCC analyses, we observed an improvement by 18.5% F1 at the patch level after first removing tiles that do not contain tumor and up to 19.4% F1 when including non-tumor tiles (Table 2) in the task of identifying whether a tile comes from a slide of grade 2 or 4 tumor. This performance gain was obtained at the expense of up to 79.6% loss in coverage.

**Table 2:**
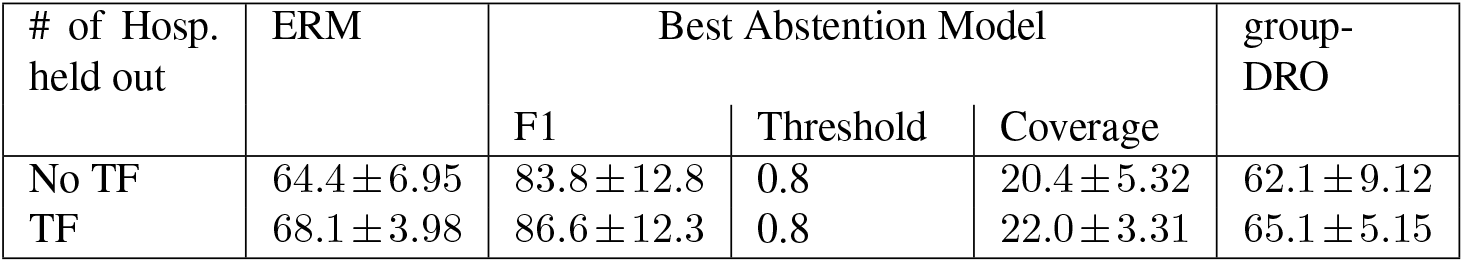
Comparing performance of proposed models in the case of identifying tumor tissue of grade II vs. grade IV in ccRCC

### 4.5. Predicting Gleason score in TCGA-PRAD

Finally, we compared the performance of a DRO and group-DRO method to a model trained with ERM on predicting Gleason score in PRAD. A model trained with group-DRO, performed comparably to a model trained with ERM. A model trained with abstention, outperformed a model trained with ERM, by up to 24.3% in grading the tumor tiles and 16.7% when all tiles are used (Tables 3 and 4). This performance boost came at 49% loss in coverage when a tumor filter was used, and 78.5% loss in coverage when no tumor filter was used.

**Table 3:** Performance of models trained to classify Gleason score of PRAD tiles as either low or high without a tumor filter

| # of Hosp.<br>held out | ERM | Best Abstention Model |  |  | group-DRO |
| --- | --- | --- | --- | --- | --- |
|  |  | F1 | Threshold | Coverage |  |
| 1 | 65.2±11.2 | 85.0±26.4 | 0.8 | 50.3±24.2 | 50.3±24.2 |
| 2 | 63.0±3.78 | 77.0±14.6 | 0.8 | 43.2±7.47 | 64.8±5.13 |
| 3 | 60.1±7.31 | 76.8±5.90 | 0.9 | 21.5±10.1 | 63.0±7.14 |
| 4 | 61.1±7.42 | 71.8±6.32 | 0.8 | 50.6±7.34 | 59.2±5.00 |
| 5 | 61.5±5.61 | 66.6±6.81 | 0.9 | 28.6±8.30 | 64.7±5.81 |

**Table 4:**
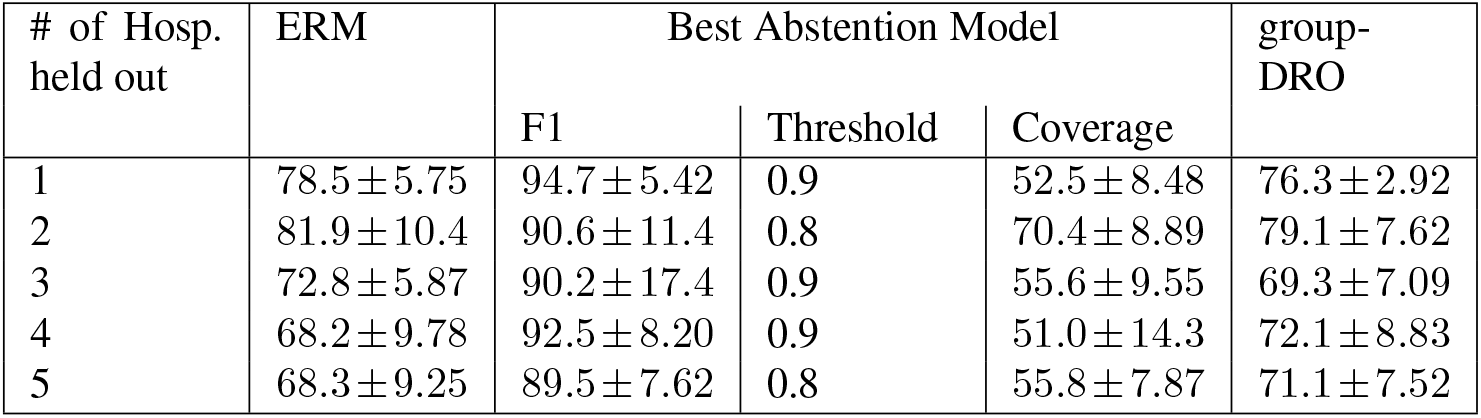
Performance of models trained to classify Gleason score of PRAD tiles as either low or high with a tumor filter

| # of Hosp.<br>held out | ERM | Best Abstention Model |  |  | group-DRO |
| --- | --- | --- | --- | --- | --- |
|  |  | F1 | Threshold | Coverage |  |
| 1 | 78.5±5.75 | 94.7±5.42 | 0.9 | 52.5±8.48 | 76.3±2.92 |
| 2 | 81.9±10.4 | 90.6±11.4 | 0.8 | 70.4±8.89 | 79.1±7.62 |
| 3 | 72.8±5.87 | 90.2±17.4 | 0.9 | 55.6±9.55 | 69.3±7.09 |
| 4 | 68.2±9.78 | 92.5±8.20 | 0.9 | 51.0±14.3 | 72.1±8.83 |
| 5 | 68.3±9.25 | 89.5±7.62 | 0.8 | 55.8±7.87 | 71.1±7.52 |

## 5. Discussion

In this study, we showed that stain profile can be used to identify the source site of a histopathology scan and contribute to significant heterogeneity in model performance. This artifact might lead a model to overfit spuriously correlated features of the slide while training on a label with weak morphological evidence.

In our analyses, we took five slides from each hospital and one hundred tiles from each slide. The differences between source sites could reflect differences specific to those tiles that were selected. However, the models’ ability to correctly identify the source site of a tile among ten sources despite using image re-coloring techniques and stain normalization implies that there are features of an image that provide sufficient visual evidence for a model to identify the source site of an image. It is possible that these features could be biological, (e.g., differences in grade, tumorinfiltrating lymphocyte infiltration, metastatic potential, or other features that are enriched in the source site’s data), so consideration of such batch effects are key for successful analysis of these data types.

We found that models achieved different performances in the task of identifying LUAD when trained on data from one hospital and tested on those of another site. We attributed this finding to a difference in the distributions of spurious variables between the training and validation datasets. We hypothesize that if a model tested well on data from a hospital while using data from other hospitals to train, the testing data are a narrow distribution of spurious and core variables that fall within the training data manifold.

We hypothesize that this approach’s capability to abstain on parts of the tissue, allows the practitioner to better understand what the model is learning from and thereby develop greater confidence in the model, as its performance relies on areas of high confidence. Ultimately, we found that DRO methods that aim to either optimize the model’s performance on a previously defined subgroup or a learned subgroup, defined in our case by the training samples that the model performed well on, were able to provide better performances on an external validation set. We make the assumption that examples that a model predicts with low confidence are OOD. However, this assumption needs further validation studies to confirm.

## 6. Conclusion

Learning spurious correlates may interfere with using models to perform biologically relevant prediction tasks and impede efforts to deliver translational care and clinical support through artificial intelligence. Machine learning applied to data from publicly available cohorts, such as the Cancer Genome Atlas (TCGA), can learn spurious correlates while trying to analyze large amounts of digitized pathology data paired with molecular and clinical outcomes, impeding multi-hospital analyses from pan-cancer patient cohorts.

Here, we evaluated the impact of batch effects and developed approaches to mitigate these fundamental challenges to digital pathology. We assessed how source sites can be learned by models, evaluated existing approaches to address known sources of batch effects, and highlighted batch effect features that, although unseen, can still impact downstream analyses. We also evaluated the role of the interpretability tool, Grad-CAM, and proposed a neural network that is robust to the distributional shifts between training and heldout test sets. Prospectively, careful consideration of seen and unseen batch effects in CV digital pathology analysis will guide successful biological investigations with potential clinical impact.

## Acknowledgements

SNH thanks Sneha Jha, Chris Labaki, Brendan Reardon and Parimarjan Negi for their helpful comments. This work was supported by R37CA222574, R01CA2227388, P50CA101942 and Dunkin’ Donuts Breakthrough Grant.

## Appendix

**Figure 5:**
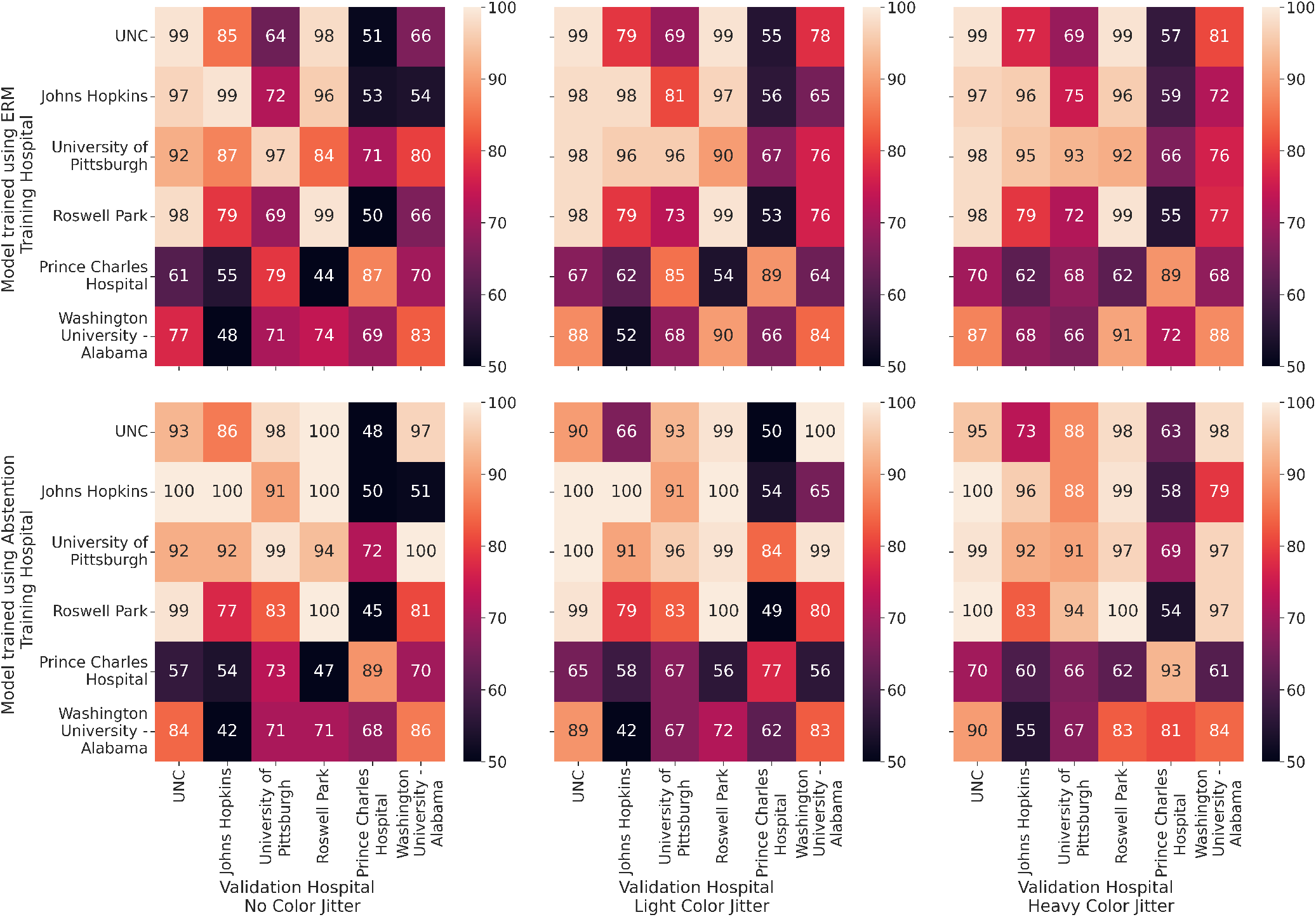
Best Validation F1 achieved by a regular CNN model and a model trained with abstention trained on one hospital (y axis) and validated on another (x axis).

**Figure 6:**
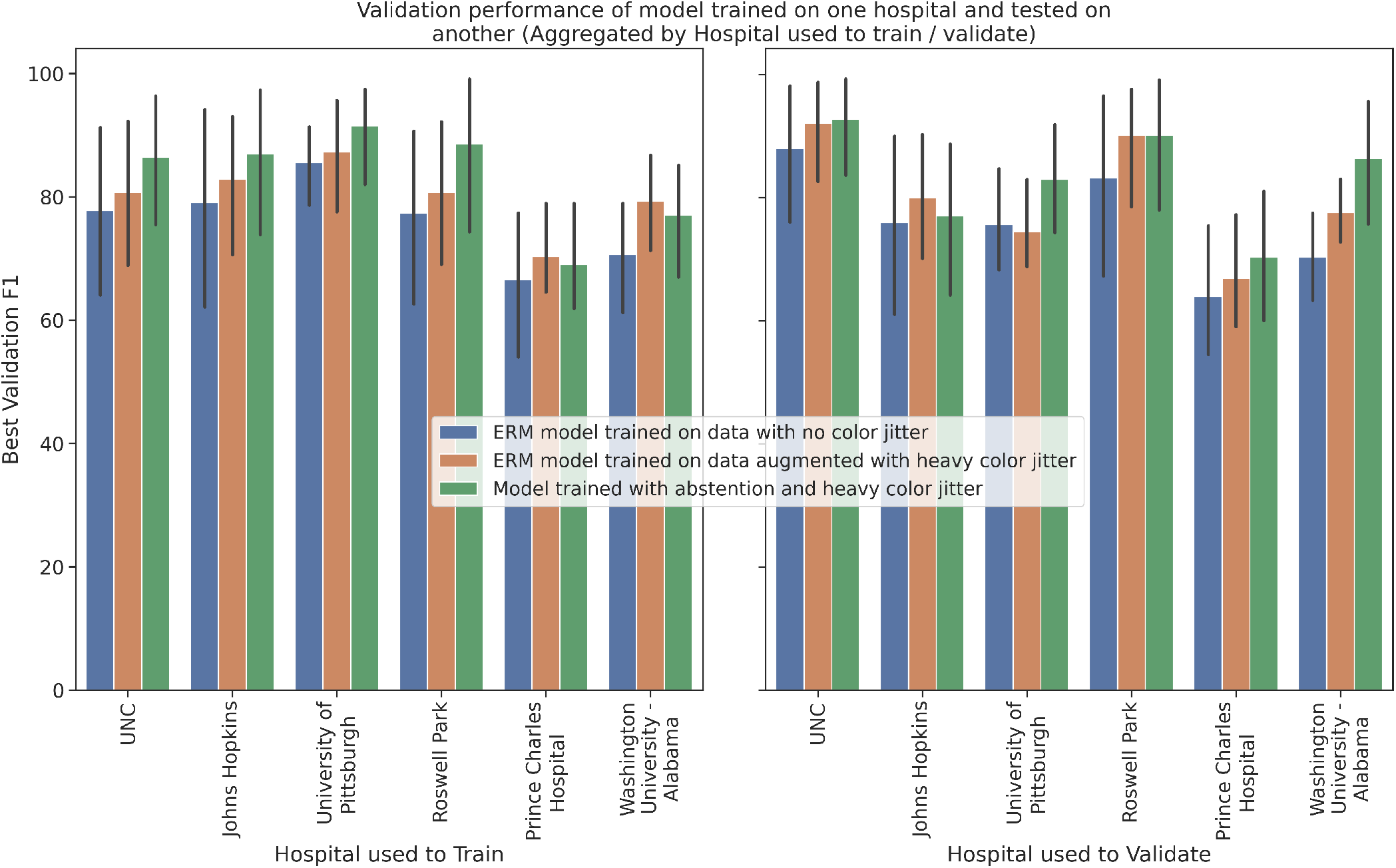
Data from subsets of Figure 5 aggregated across hospitals used to train and validate

**Figure 7:**
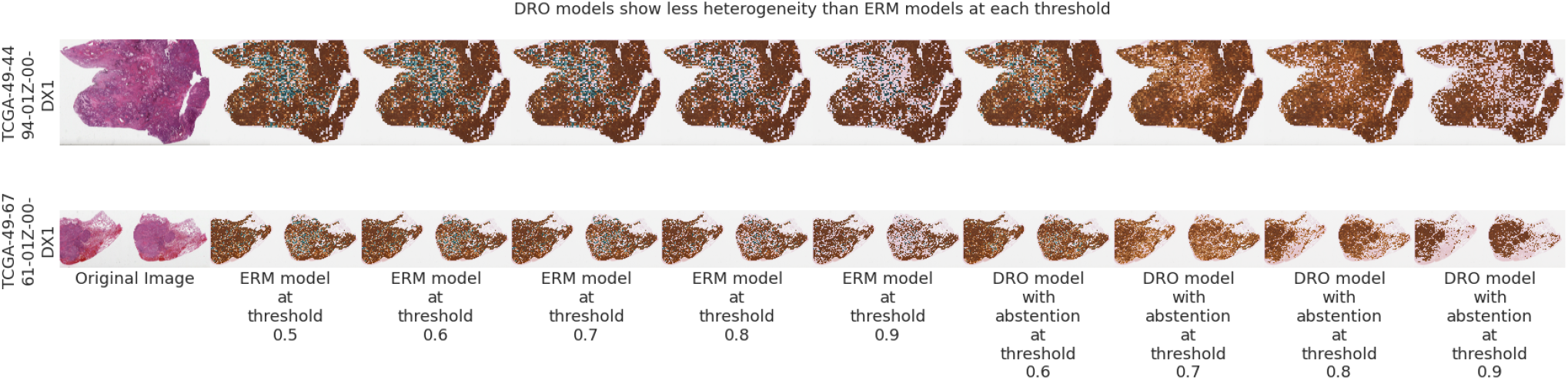
Reduced heterogeneity in a model trained using group-DRO for tumor versus normal identification in LUAD. Brown indicates patches that were predicted as tumor, blue indicates patches that were predicted as normal, surrounding tissue. Models trained with more stringent abstention show more uniform class prediction by abstaining on tiles where the features on the tile are out-of-distribution relative to the features pertinent to the WSI label (first row). Second row: group-DRO methods abstain on tiles on the right hand side of the tissue where the tissue does not bear tumor, as verified by an expert. ERM methods call non-tumor region as tumor, even at high confidence thresholds.

**Figure 8:**
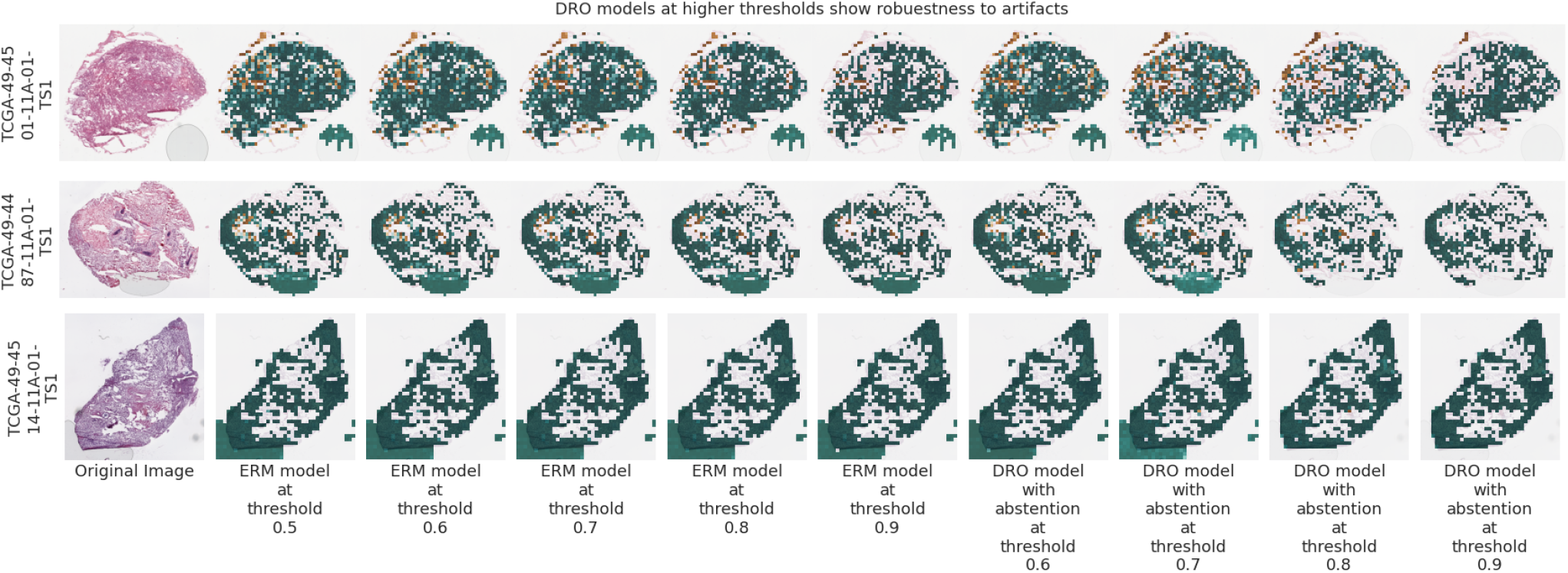
ERM methods predict bubble artifacts as healthy surrounding tissue. Models trained with stringent abstention abstain from making predictions on artifacts.

**Table 5:** Comparing performance of proposed models in the task of identifying tumor tissue in LUAD

| # Hosp.<br>held out | ERM | Abstention threshold |  |  |  | group-DRO |
| --- | --- | --- | --- | --- | --- | --- |
|  |  | 0.6 | 0.7 | 0.8 | 0.9 |  |
| 1 | 91.1±4.05 | 93.1±4.09 | 96.6±8.35 | 98.4±7.89 | 97.3±4.84 | 92.2± 8.08 |
| 2 | 88.3±2.29 | 88.6±1.50 | 89.1±3.63 | 93.3±3.94 | 95.2±4.39 | 88.6± 3.99 |
| 3 | 78.6±8.78 | 77.5±7.75 | 78.9±7.71 | 78.6±9.50 | 82.0±9.26 | 83.6± 9.84 |
| 4 | 81.1±8.86 | 78.9±9.96 | 80.4±19.4 | 84.0±8.89 | 91.0±7.70 | 88.8± 5.85 |
| 5 | 72.2±3.84 | 73.4±3.88 | 71.9±4.40 | 76.3±3.61 | 79.7±7.37 | 77.4± 5.09 |

**Table 6:** Comparing coverage of proposed models in the task of identifying tumor tissue in LUAD

| # of Hosp.<br>held out | Abstention threshold |  |  |  |
| --- | --- | --- | --- | --- |
|  | 0.6 | 0.7 | 0.8 | 0.9 |
| 1 | 98.1±1.14 | 94.5±8.80 | 86.6±5.80 | 63.2±21.0 |
| 2 | 94.6±1.60 | 87.4±5.63 | 79.1±10.9 | 54.3±15.0 |
| 3 | 95.3±1.51 | 87.3±8.15 | 83.7±12.4 | 65.2±20.8 |
| 4 | 94.5±3.08 | 84.9±28.2 | 74.0±24.1 | 54.8±21.8 |
| 5 | 95.5±3.64 | 90.4±13.7 | 83.6±16.4 | 59.6±23.8 |

**Table 7:** Comparing performance of proposed models in the case of identifying tumor tissue of grade II vs. grade IV in ccRCC

|  | ERM | Abstention threshold |  |  |  | group-DRO |
| --- | --- | --- | --- | --- | --- | --- |
|  |  | 0.6 | 0.7 | 0.8 | 0.9 |  |
| No TF | 64.4±6.95 | 68.6±8.83 | 71.2±14.6 | 83.8±12.8 | 76.1±15.1 | 62.1±9.12 |
| TF | 68.1± 3.98 | 69.7±10.5 | 72.3±10.9 | 86.6±12.3 | 73.2±16.0 | 65.1±5.15 |

**Table 8:** Comparing coverage of proposed models in the case of identifying tumor tissue of grade II vs. grade IV in ccRCC

|  | Abstention threshold |  |  |  |
| --- | --- | --- | --- | --- |
|  | 0.6 | 0.7 | 0.8 | 0.9 |
| No TF | 65.4±19.8 | 37.3±17.3 | 20.4±5.32 | 8.07±10.3 |
| TF | 78.7±13.0 | 30.4±11.7 | 22.0±3.31 | 6.03±7.74 |

**Table 9:**
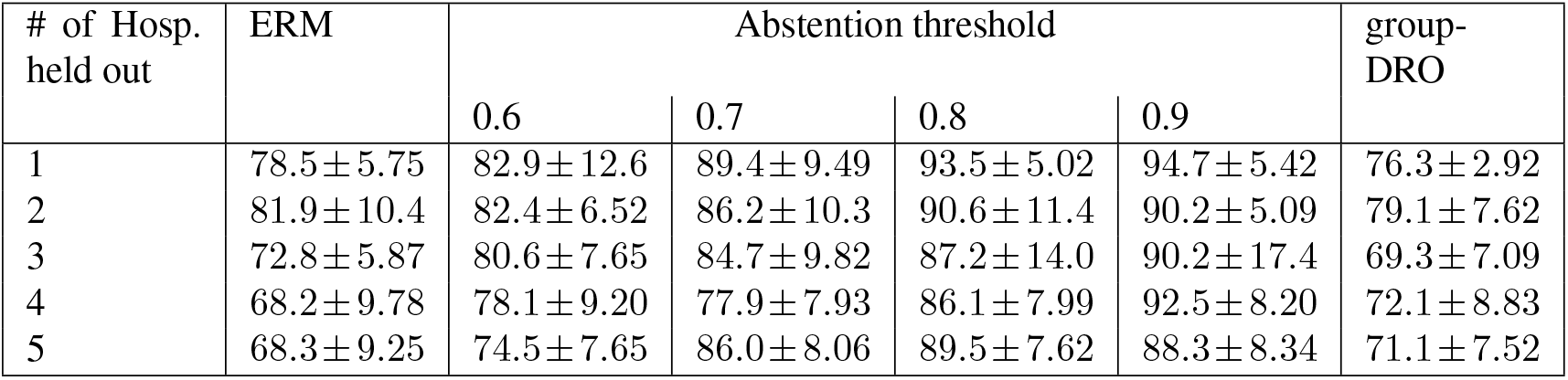
Comparing performance of models trained to classify Gleason score of PRAD tiles as either low or high with a tumor filter

**Table 10:** Comparing coverage of models trained to classify Gleason score of PRAD tiles as either low or high with a tumor filter

| # of Hosp.<br>held out | Abstention threshold |  |  |  |
| --- | --- | --- | --- | --- |
|  | 0.6 | 0.7 | 0.8 | 0.9 |
| 1 | 88.4 ± 4.21 | 80.6 ± 2.57 | 69.5 ± 5.05 | 52.5 ± 8.48 |
| 2 | 89.5 ± 3.43 | 81.4 ± 7.64 | 70.4 ± 8.89 | 46.3 ± 16.8 |
| 3 | 92.0 ± 1.42 | 80.7 ± 4.05 | 70.3 ± 4.36 | 55.6 ± 9.55 |
| 4 | 90.2 ± 3.45 | 77.4 ± 5.11 | 65.7 ± 6.87 | 51.0 ± 14.3 |
| 5 | 89.4 ± 1.22 | 74.7 ± 9.82 | 55.8 ± 7.87 | 39.0 ± 7.18 |

**Table 11:**
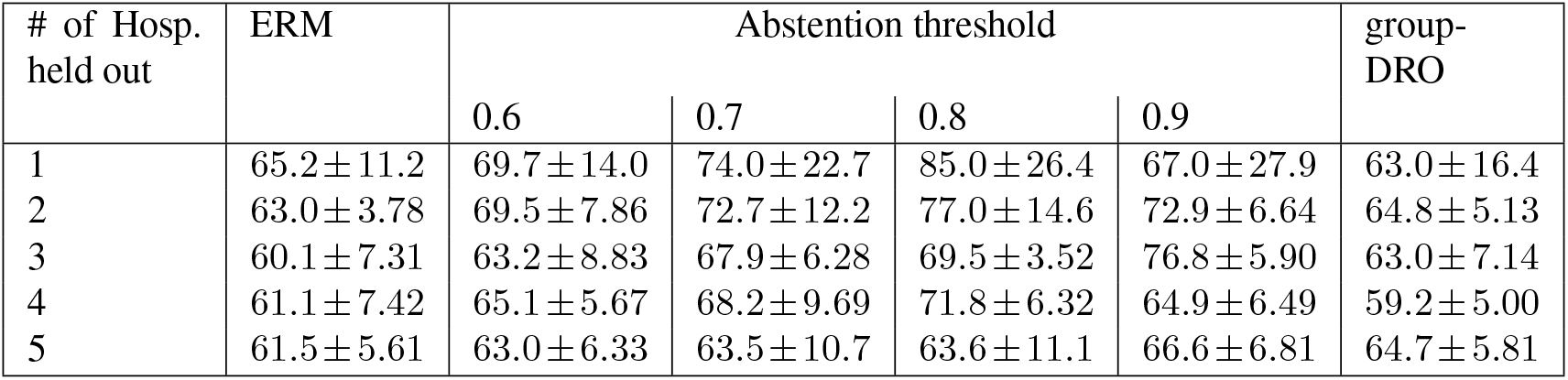
Comparing performance of models trained to classify Gleason score of PRAD tiles as either low or high without a tumor filter

| # of Hosp.<br>held out | ERM | Abstention threshold |  |  |  | group-DRO |
| --- | --- | --- | --- | --- | --- | --- |
|  |  | 0.6 | 0.7 | 0.8 | 0.9 |  |
| 1 | 65.2 ± 11.2 | 69.7 ± 14.0 | 74.0 ± 22.7 | 85.0 ± 26.4 | 67.0 ± 27.9 | 63.0 ± 16.4 |
| 2 | 63.0 ± 3.78 | 69.5 ± 7.86 | 72.7 ± 12.2 | 77.0 ± 14.6 | 72.9 ± 6.64 | 64.8 ± 5.13 |
| 3 | 60.1 ± 7.31 | 63.2 ± 8.83 | 67.9 ± 6.28 | 69.5 ± 3.52 | 76.8 ± 5.90 | 63.0 ± 7.14 |
| 4 | 61.1 ± 7.42 | 65.1 ± 5.67 | 68.2 ± 9.69 | 71.8 ± 6.32 | 64.9 ± 6.49 | 59.2 ± 5.00 |
| 5 | 61.5 ± 5.61 | 63.0 ± 6.33 | 63.5 ± 10.7 | 63.6 ± 11.1 | 66.6 ± 6.81 | 64.7 ± 5.81 |

**Table 12:** Comparing coverage of models trained to classify Gleason score of PRAD tiles as either low or high without a tumor filter

| # of Hosp.<br>held out | Abstention threshold |  |  |  |
| --- | --- | --- | --- | --- |
|  | 0.6 | 0.7 | 0.8 | 0.9 |
| 1 | 83.5 ± 3.15 | 63.5 ± 8.19 | 50.3 ± 24.2 | 37.4 ± 30.4 |
| 2 | 84.8 ± 4.49 | 70.6 ± 4.01 | 43.2 ± 7.47 | 22.4 ± 10.4 |
| 3 | 86.7 ± 5.40 | 62.1 ± 5.90 | 51.2 ± 9.91 | 21.5 ± 10.1 |
| 4 | 87.5 ± 2.38 | 65.6 ± 8.61 | 50.6 ± 7.34 | 31.6 ± 6.47 |
| 5 | 86.5 ± 5.37 | 65.4 ± 7.58 | 47.8 ± 8.73 | 28.6 ± 8.30 |

